# From Fragmentation to Resolution: High-Fidelity Genome Assembly of *Zancudomyces culisetae* through Comparative Insights from PacBio, Nanopore, and Illumina Sequencing

**DOI:** 10.1101/2025.04.24.650499

**Authors:** Huimei Yang, Yan Wang

**Author notes:** **Corresponding author**: Yan Wang.

## Abstract

*Zancudomyces culisetae* is an obligate symbiotic fungus inhabiting the digestive tracts of aquatic insect larvae, including black flies, midges, and mosquitoes. With a global distribution and high prevalence in disease-transmitting insects, *Z. culisetae* serves as a model for studying insect gut fungi. A previous draft genome assembly using Illumina short reads provided insights into its genome composition, such as a low GC ratio and evidence of horizontal gene transfer. However, its fragmented nature has limited deeper exploration of the evolutionary mechanisms shaping these gut symbionts. To address this gap, we generated a wealth of genomic resources for *Z. culisetae* using multiple sequencing platforms, including Illumina, Oxford Nanopore, PacBio-CLR (Complete Long Reads), and PacBio-HiFi (High Fidelity). This also provides an opportunity to compare these popular sequencing methods to suggest the optimal approach for fungal genome assembly. Our results suggest that PacBio-HiFi produced the most complete assembly, yielding a 27.8 Mb genome size with 26 contigs, representing the highest-quality genome of insect gut fungi to date. Additionally, we generated transcriptomic data to support genome annotation, identifying 8,484 protein-coding genes. Despite the improved genome quality, *Z. culisetae* lacks approximately 20% of Benchmarking Universal Single-Copy Orthologue (BUSCO) commonly found in fungi, reflecting adaptations to its obligate symbiotic lifestyle. This study not only provides valuable genomic resources for insect gut fungal research but also evaluates the strengths and limitations of current genome sequencing and assembly approaches, offering best practices for fungal genome analysis and genetic research.

**Article Summary:** Mosquito larvae harbor gut-dwelling fungi that may influence their development and disease transmission. *Zancudomyces culisetae* is a widespread fungal symbiont of mosquitoes, but its genome remained incomplete. This study explored and compared popular sequencing technologies, including Illumina, Oxford Nanopore, PacBio-CLR, and PacBio-HiFi, to generate a high-quality genome, alongside the first transcriptomic data. PacBio-HiFi produced the most complete assembly (27.8 Mb, 26 contigs) with 8,484 protein-coding genes, though ∼20% of core fungal genes were absent, reflecting symbiotic adaptation. This study establishes a genomic foundation for insect gut-dwelling fungi and evaluates available sequencing and assembly strategies for high-quality fungal genome production.

## Introduction

*Zancudomyces culisetae* (formerly known as *Smittium culisetae*) is a well-studied insect gut-dwelling fungus, belonging to an important lineage of early-diverging fungi (Harpellales, Zoopagomycota (White 2006; Wang et al. 2013; Spatafora et al. 2016). *Zancudomyces culisetae* has been frequently reported worldwide, primarily colonizing the hindgut of disease-vector insect larvae, including black flies, mosquitoes, and midges (Lichtwardt 1986; Lichtwardt et al. 1999; Valle and Santamaria 2004; White et al. 2006; Chaudhary et al. 2023). These fungi have maintained an obligate symbiotic relationship with dipteran hosts for approximately 270 million years, coinciding with the evolution of complete metamorphosis in insects (Wang et al. 2019). This ancient symbiosis offers a unique model for studying microbial adaptation to extreme environments, such as insect guts, and the long-term co-evolution between fungi and one of Earth’s most ecologically dominant animal groups.

The first *Z. culisetae* culture was isolated from a fourth-instar larva of *Culiseta incidens* and successfully grown on SNB-9 medium and Difco brain-heart infusion agar (Clark et al. 1963). Using similar techniques, additional axenic cultures were established from various location (Lichtwardt 1964; Lichtwardt and Williams 1990; Strongman and White 2008). These cultures enabled extensive research into the biology and symbiotic interactions of insect gut-dwelling fungi, including their lifecycle, nutritional requirements, and host distribution (Lichtwardt 1964; Horn and Lichtwardt 1981; Williams 1983; Horn 1989; Sato 1992; McCreadie and Beard 2003; Nelder et al. 2005; Vojvodic and McCreadie 2009). More recently, molecular systematics have supported the independent placement of *Z. culisetae* outside *Smittium*, where it was originally classified (Wang et al. 2013; Tretter et al. 2014; Wang et al. 2014). The successful cultivation of *Z. culisetae* facilitated the whole genome sequencing using Illumina short reads, leading to the discovery of a horizontally transferred polyubiquitin gene from mosquito hosts (Wang et al. 2016). Subsequent genomic studies on insect gut fungi have uncovered a noncanonical genetic mechanism in which the UGA stop codon is repurposed to encode the 21^st^ amino acid, selenocysteine, in fungi for the first time (Mariotti et al. 2019). However, the currently available *Z. culisetae* genome assembly (COL-18-3; ARSEF 9012) was generated using Illumina short-read sequencing, resulting in a highly fragmented assembly of 1,954 scaffolds (Wang et al. 2016). This fragmentation limits further investigation into critical molecular mechanisms, such as horizontal gene transfer and selenocysteine incorporation via stop codon reassignment. Additionally, no RNA sequencing data are available for accurate gene annotation or expression analysis in *Z. culisetae*.

To address these limitations, we present a reference-quality genome assembly of *Z. culisetae* in this study. We sequenced the HAW-14-8 strain (ARSEF 9014), originally isolated from the hindgut of an *Aedes albopictus* mosquito in Oahu (Hawaii, USA). A combination of cutting-edge sequencing technologies, including PacBio Single-Molecule Real-Time (SMRT) Sequel II platform (for both continuous long reads and circular consensus sequencing reads, also known as High-Fidelity reads), Oxford Nanopore PromethION, and Illumina NovaSeq PE150, allowed us to compare the performance of different sequencing platforms and assembly strategies for gut-dwelling fungi. The reference-quality genome assembly exhibited relatively low completeness scores indicated by the Benchmarking Universal Single-Copy Orthologue (BUSCO), which is consistent with findings from previously sequenced gut-dwelling fungal genomes. Our study provides a high-quality genomic resource that will facilitate further investigations into the evolutionary adaptation, molecular interactions, and functional genomics of insect gut fungi, including possible gene loss events due to symbiotic interactions with mosquito hosts.

## Materials and Methods

### Fungal sample preparation and DNA extraction

The *Zancudomyces culisetae* strain HAW-14-8 was originally isolated from *Aedes albopictus* in Oahu, Hawaii, USA (Lichtwardt 1964) and obtained from the USDA-ARS Collection of Entomopathogenic Fungal Cultures (ARSEF 9014). Fungal cultures were grown in Brain Heart Infusion Glucose Tryptone vitamins (BHIGTv) broth at room temperature for one week, following the method described by White et al. (2006). Fungal tissues were harvested by filtration through autoclaved Miracloth (MilliporeSigma, Canada) and ground to a fine powder in liquid nitrogen. High molecular weight DNA (>20kb) was extracted using the MagAttract HMW DNA Kit (QIAGEN, Canada), while mRNA was isolated using the MagMax plant RNA isolation kit (Thermo Fisher). Both DNA and RNA extractions followed the instructions provided by the respective manufacturers.

### Genome sequencing

DNA and RNA libraries (PE150 for 350bp) were prepared and sequenced on the Illumina NovaSeq 6000 system at the Novogene UC Davis Sequencing Center. Long-read DNA libraries (>20kb) were prepared and sequenced at the Centre for Applied Genomics, Hospital for Sick Children (Toronto, Canada) using both the ONT PromethION (FLO-PRO002 R9.4 flow cell with SQK-LSK110 ligation sequencing kit) and PacBio Sequel II platforms (Express TPK 2.0 + Sequencing primer v4) separately.

### Genome assembly

Illumina short reads were adapter-trimmed using Trimmomatic v0.36 (Bolger et al. 2014) and quality-checked using FASTQC v0.11.9 before being assembled into contigs using SPAdes v3.13.1 with default k-mer setting for 150bp read length. Raw electrical signal FAST5 files from the ONT PromethION platform were base-called using Guppy version 5.0.12 (https://community.nanoporetech.com) and the resulting reads were assembled into contigs with Canu v2.2 (Zimin et al. 2017). The PacBio CLR raw reads were assembled using Flye (Kolmogorov et al. 2019), while PacBio HiFi raw data were adapter-trimmed with HiFiAdapterFilt v2.0.0 and assembled into contigs using HiFiASM v0.16.1-r375 (Cheng et al. 2021; Sim et al. 2022). FinisherSC was applied to improve the assembly by refining the HiFiASM primary assembly using the trimmed long-read data (Lam et al. 2015). For hybrid assemblies, SPAdes v3.13.1 was used to integrate Illumina short-reads with long-reads from PacBio CLR and/or Oxford Nanopore, leveraging long-read data for gap closure and repeat resolution (Antipov et al. 2016). An overview of the workflow and comparison schemes is illustrated in Figure 1.

**Figure 1.**
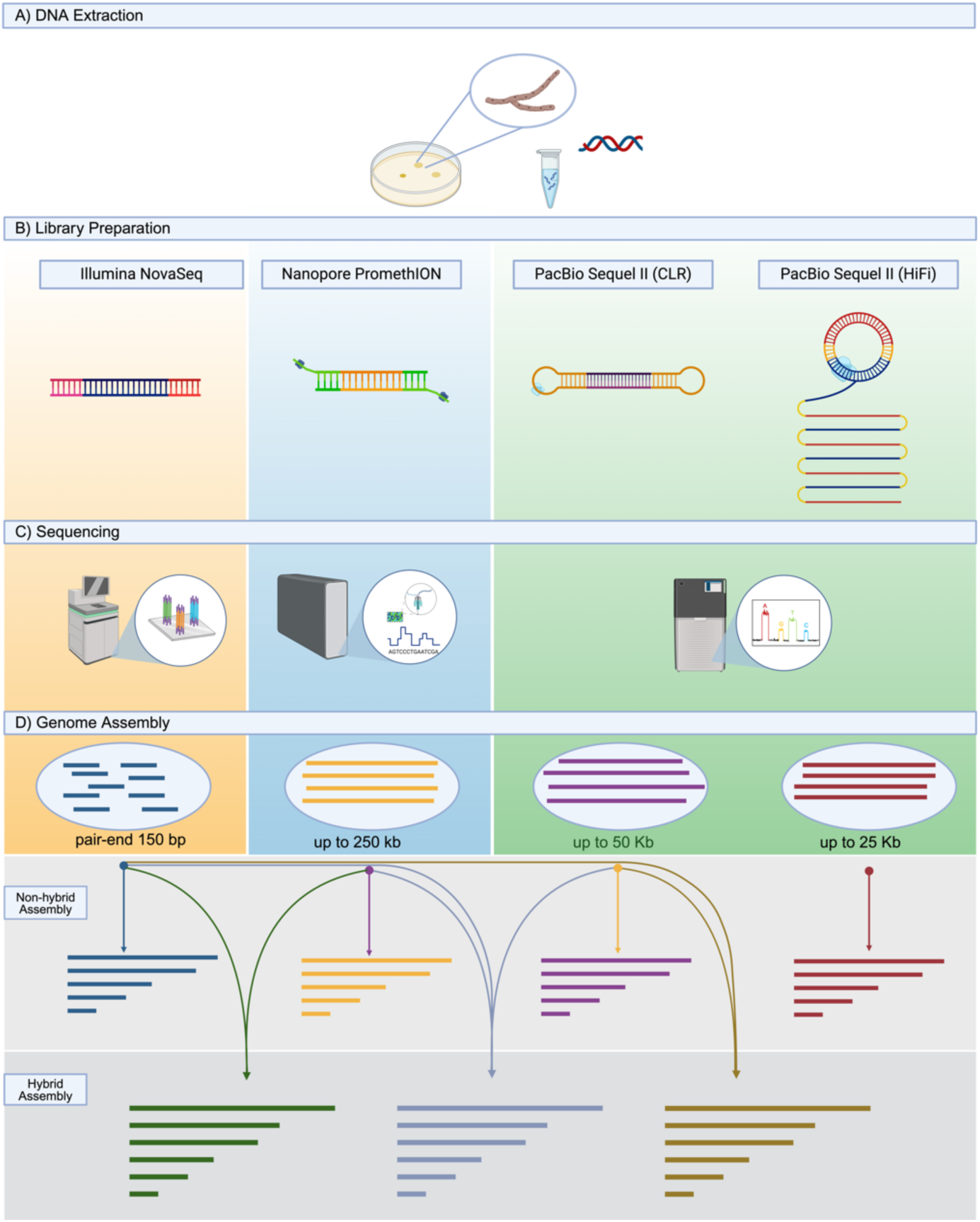
Schematic overview of the sequencing pipeline and assembly strategies using Illumina NovaSeq, Oxford Nanopore Technologies PromethION (Nanopore), Pacific Biosciences Sequel II Continuous Long Read (PacBio-CLR), and Pacific Biosciences Sequel II High Fidelity Long Read (PacBio-HiFi). The workflow includes DNA extraction, library preparation, sequencing, and genome assembly. (A) DNA Extraction: High-molecular-weight DNA was extracted from fungal cultures for all sequencing platforms. (B) Library Preparation: DNA fragments of platform-specific sizes were prepared, followed by adapter ligation. (C) Sequencing: Raw reads were generated as either short-reads (Illumina NovaSeq) or long-reads (Nanopore, PacBio-CLR, and PacBio-HiFi). (D) Genome Assembly: Seven genome assemblies were produced using either non-hybrid strategies (using single-platform reads) or hybrid strategies (combining reads from multiple platforms).

### Assessment and comparative analysis of genome assemblies

Genome assembly completeness was assessed using Benchmarking Universal Single-Copy Orthologs (BUSCO) v5.1.2 with the fungi_odb10 database (Manni et al. 2021). Genome assembly statistics were summarized using gfastats v1.3.10 and visualized using the R package “ggplot2” and BlobToolKit v2.6.5 (Wickham 2016; Challis et al. 2020; Formenti et al. 2022). The assemblies, generated using different strategies, were first reordered against the highest-quality reference (i.e., PacBio-HiFi) using Mauve Contig Mover (MCM) tool, followed by Locally Collinear Blocks (LCBs) identification via progressiveMauve alignment with default settings (Rissman et al. 2009; Darling et al. 2010).

### Genome annotation

Contigs shorter than 1kb were removed using BioKIT before proceeding with annotation (Steenwyk et al. 2022). RepeatModeler v2.0.2a and Repeatmasker v4.1.0 (https://www.repeatmasker.org/RepeatModeler/) were used to softmask interspersed repeats and low-complexity DNA sequences in the assembly. RNAseq raw data were adapter-trimmed using Trimmomatic v0.36 and mapped to the genome assembly using HISAT2 v2.2.1, followed by processing with SAMtools v1.12. Genome annotation was performed on the softmasked genome assembly using the Braker pipeline v1.9 (parameters –fungus –softmasking 1), which integrates Augustus and GeneMark with RNA evidence for gene prediction. Function annotations were conducted using PfamScan v1.6 to identify protein domains by searching the predicted *Z. culisetae* proteins against the Pfam v35.0 HMM database (Finn et al. 2016).

## Results

Genome assemblies generated using different sequencing platforms and assembly strategies were evaluated for completeness using BUSCO (Benchmarking Universal Single-Copy Orthologs) (Figure 2A). Most assemblies exhibited similar completeness scores, ranging from 70.1% to 71.9% for the Complete category, which includes both Single Copy and Duplicated content. The exception was the assembly generated from Nanopore data alone, which had a lower completeness score of 63.7%. Notably, while Illumina-only and hybrid approaches showed comparable completeness to long-read assemblies, the Nanopore and PacBio-HiFi assemblies displayed a slightly higher proportion of Complete-Duplicated BUSCO content.

**Figure 2.**
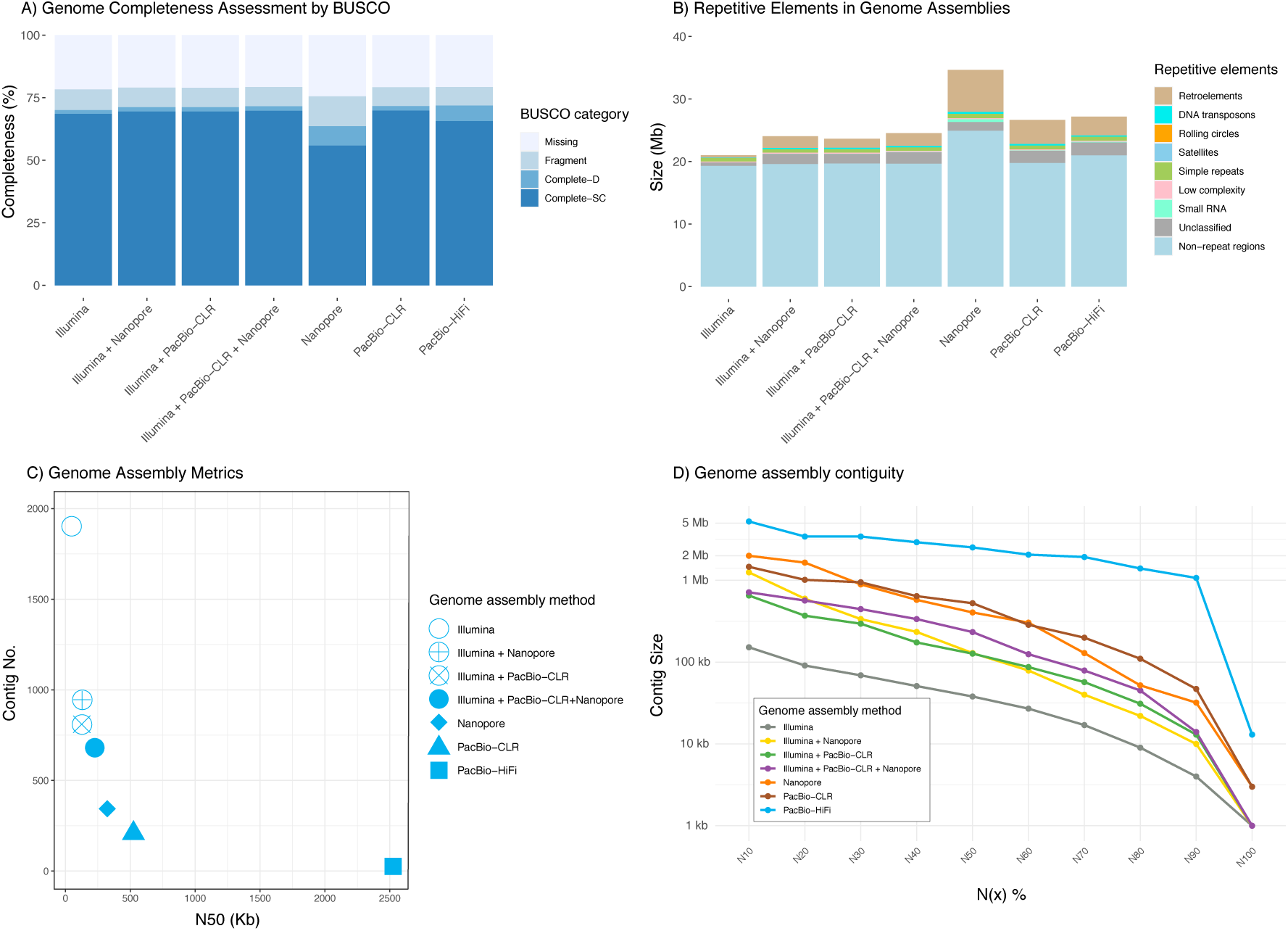
Comparison of genome sequencing and assembly approaches for *Z. culisetae*. A) Genome assembly completeness assessed using BUSCO for seven assemblies generated from single-platform (Illumina, Nanopore, PacBio-CLR, or PacBio-HiFi) or hybrid approaches (Illumina + Nanopore, Illumina + PacBio-CLR, and Illumina + PacBio-CLR + Nanopore). Stacked bars represent the proportion of complete (single-copy/duplicated), fragmented, and missing BUSCO genes. B) Repetitive element content annotated with RepeatMasker. Stacked bars show the relative abundance of repeat classes (e.g., Retroelements, DNA transposons, Simple repeats) in each assembly. C) Genome assembly metrics plot comparing N50 (x-axis) and total contig count (y-axis). Symbols denote assembly strategies. D) Assembly contiguity profiles plotting contig sizes (y-axis) against N(x) statistics (x-axis), where N(x) indicates the length for which the collection of all contigs of that length or longer covering x% of the genome (e.g., N50, N90).

Repetitive elements were analyzed and compared across genome assemblies generated using different approaches (Figure 2B). The total repetitive content varied among assemblies, with long-read assemblies (e.g., Nanopore, PacBio-CLR, and PacBio-HiFi) recovering a larger proportion of repetitive sequences compared to Illumina-only assemblies. Hybrid assembly approaches showed significantly improvement by recovering a greater number of repetitive elements than short-read assembly alone. Interestingly, Nanopore assembly reports the largest number of repetitive elements, contributing to its larger genome size. Among the repeat categories, retroelements constituted the majority of the repetitive content across all assemblies, followed by unclassified repeats, simple repeats, and DNA transposons. Notably, the Nanopore-based assembly recovered the largest region of repetitive elements in nearly every category, including retroelements, DNA transposons, small RNA, simple repeats, and low complexity.

Satellites and rolling circles were not consistently recovered across assemblies that the rolling circles were absent in Illumina-only and Nanopore-only assemblies, while satellites were only detected in hybrid assemblies (Illumina + PacBio-CLR and Illumina + PacBio-CLR + Nanopore).

To evaluate the quality of genome assemblies generated by different sequencing strategies, we compared key assembly metrics, including N50 and contig counts (Figure 2C), as well as contig length statistics (Figure 2D). Assemblies produced using long-read technologies (e.g., PacBio-CLR, PacBio-HiFi, and Nanopore) consistently achieved higher N(x) values (e.g., N50 and N90; Figure 2D) and fewer contig numbers (Figure 2C), indicating improved assembly continuity.

Among all utilized methods in this study, the PacBio-HiFi assembly outperformed others, exhibiting the highest assembly statistics, including N50 (2.5 Mb) and the fewest contigs (26), reflecting superior contiguity and minimal fragmentation (Figure 2 C and D). In contrast, the assembly generated using Illumina-only reads showed the lowest N50 (38 Kb) and the largest number of contigs (1,898), indicating significant fragmentation and large number of gaps remained within the assembly. Hybrid assembly approaches, which combine Illumina short-read data with long-read data for gap closure and repeat resolution, demonstrated improved performance (Figure 2D). For example, combining Illumina with Nanopore data significantly reduced the contig number (from 1,898 to 953) and increased the N50 score (from 37,998 to 128,795) (Figure 2C). Similarly, the combination of Illumina and PacBio-CLR data resulted in fewer contigs (810 vs. 952) and a slightly lower N50 score (126,571 vs. 128,795). The metrics improved further when using both types of long-read data, yielding 688 contigs and an N50 of 232,805. These results highlight that enhanced continuity can be achieved by incorporating long-reads to Illumina short-read assemblies. Overall, assemblies based solely on long-read data generally performed better, with the peak performance achieved by PacBio-HiFi reads with 26 contigs and N50 of 2.5Mb. As such, the PacBio-HiFi assembly was used as the reference for whole-genome alignment of *Z. culisetae* assemblies generated using different sequencing platforms and combinations, as visualized with progressiveMauve (Figure 3). Most Locally Collinear Blocks (LCBs) are recovered across all assemblies, however, the interspersed sequences between LCBs are not consistently present and appear reduced in size, particularly in the Illumina-based assemblies. Notably, the unaligned regions at the far right of each assembly are most extensive in the Nanopore assembly, contributing to its overall larger genome size compared to the other assemblies.

**Figure 3.**
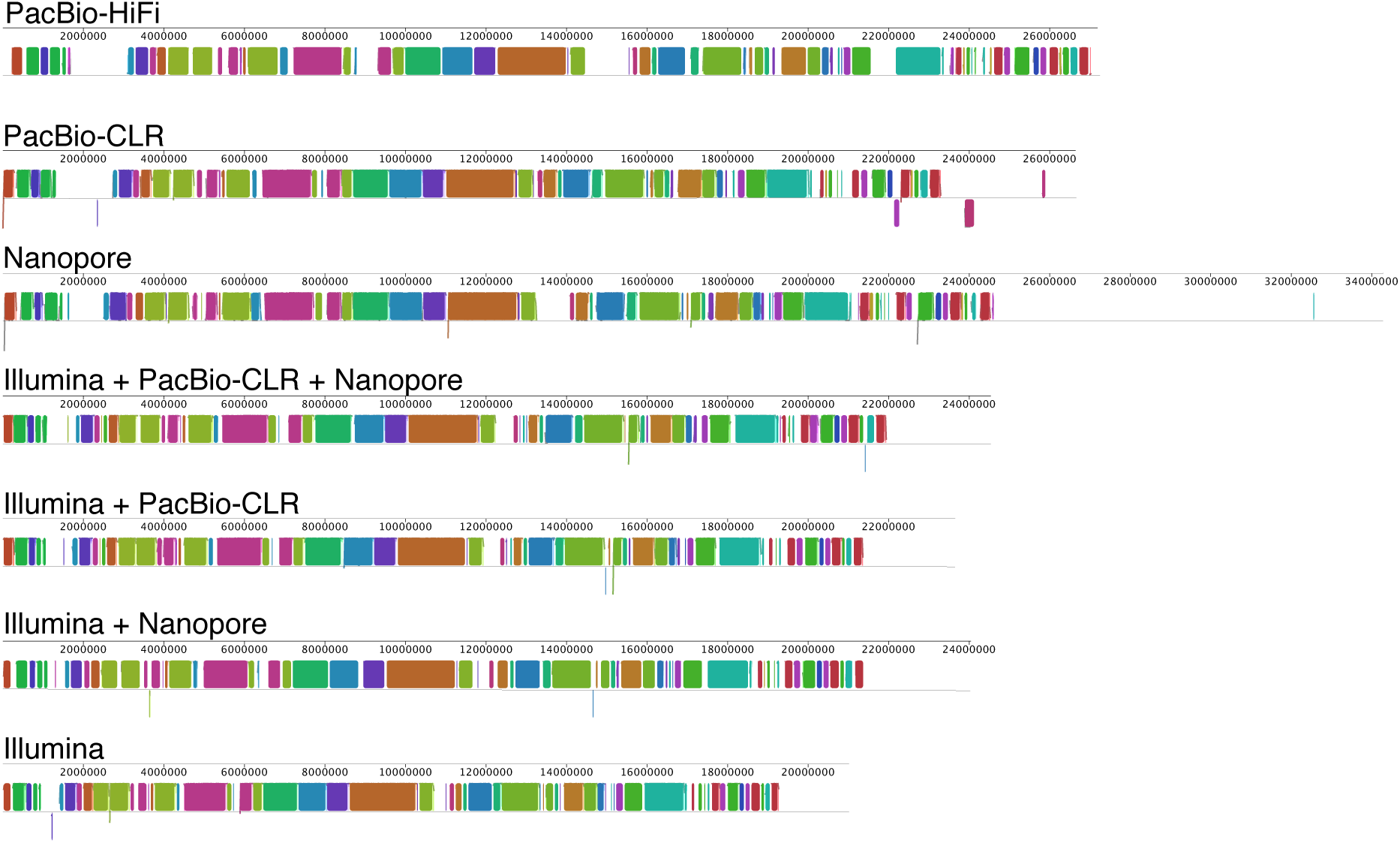
Whole-genome alignment of *Z. culisetae* assemblies using progressiveMauve. The PacBio-HiFi assembly (top row) serves as the reference for comparison with other assemblies (labeled rows below). The horizontal axis represents the size of each genome assembly. Locally Collinear Blocks (LCBs) are shown as colored regions, with shared colors indicating homologous sequences across assemblies. Inverted blocks (reverse orientation) appear below the horizontal axis. White regions indicate sequences that did not meet the minimum weight threshold set by the progressiveMauve algorithm.

The highest-quality *Z. culisetae* assembly was generated using PacBio HiFi reads, resulting in 26 contigs with a total length of 27.2 Mb and a GC content of 35.56% (Figure 4 and Table 1). The largest contig exceeds 5.2 Mb, and 11 contigs are over 1 Mb, together comprising more than 93% of the genome (Figure 4). The genome encodes 8,484 genes, with annotation predicting 10,952 Pfam domains, representing 3,026 distinct Pfam types. Transcriptome reads mapped to the genome at a rate of 97.81%, supporting its high completeness and accuracy.

**Figure 4.**
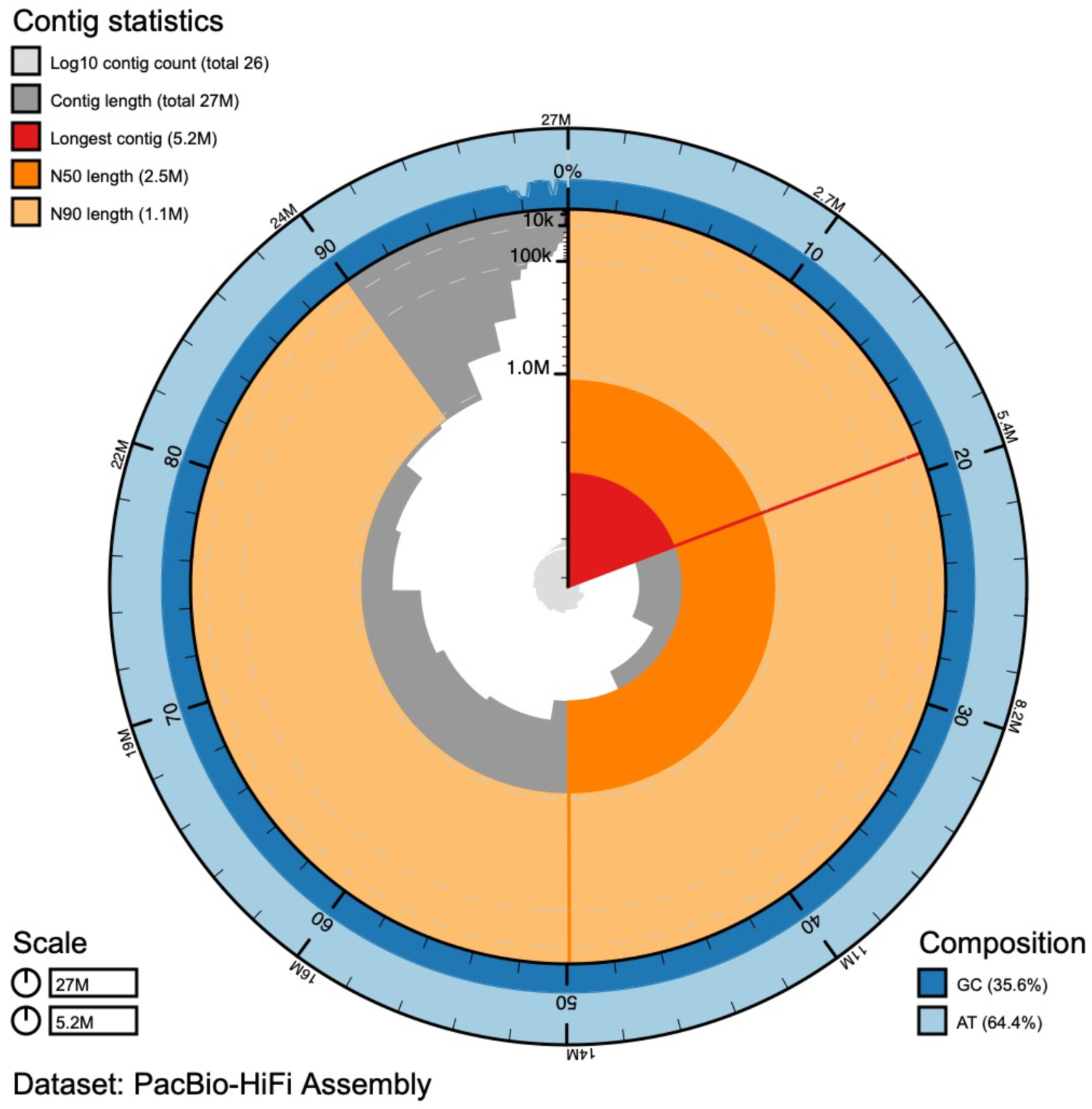
Snail plot of the best assembly for *Zancudomyces culisetae*, generated from PacBio-HiFi reads. The longest contig is highlighted in red (5.2 Mb), with remaining contigs arranged clockwise by size. Dark and light orange arcs indicate N50 and N90 values, respectively. The central gray spiral shows cumulative contig count on a log scale, with white lines marking each order of magnitude. The GC content is displayed as dark and light blue tracks.

**Table 1.** Summary of genome assembly statistics for *Zancudomyces culisetae* using different sequencing technologies.

| Sequencing method | Illumina | Illumina & Nanopore | Illumina & PacBio-CLR | Illumina & PacBio-CLR & Nanopore | Nanopore | PacBio-CLR | PacBio-HiFi |
| --- | --- | --- | --- | --- | --- | --- | --- |
| Assembler | SPAdes | SPAdes | SPAdes | SPAdes | Canu | Flye | HiFiASM & Finisher |
| Genome size (bp) | 21,012,904 | 24,039,265 | 23,647,634 | 24,535,143 | 34,666,050 | 26,640,971 | 27,184,066 |
| Contigs No. | 1,898 | 952 | 810 | 688 | 344 | 208 | 26 |
| GC ratio (%) | 35.67 | 35.47 | 35.46 | 35.42 | 35.28 | 35.58 | 35.56 |
| Largest frg | 367,220 | 1,383,685 | 1,061,931 | 1,065,925 | 3,086,553 | 1,691,663 | 5,239,357 |
| N50 | 37,998 | 128,795 | 126,571 | 232,805 | 322,842 | 525,018 | 2,524,970 |
| L50 | 139 | 32 | 41 | 28 | 24 | 15 | 4 |
| BUSCO %<br>(complete [single-copy, duplicated], fragmented, missing) | 70.1[68.6, 1.5], 8.3, 21.6 | 71.3[69.5, 1.8], 7.8, 20.9 | 71.3[69.5, 1.8], 7.7, 21.0 | 71.6[69.8, 1.8%], 7.7, 20.7 | 63.7[55.9, 7.8], 11.9, 24.4 | 71.7[69.9, 1.8], 7.5, 20.8 | 71.9[65.7, 6.2], 7.4, 20.7 |
| Retroelements (bp) | 414,226 | 1,888,948 | 1,470,941 | 2,056,224 | 6,723,120 | 3,836,832 | 3,031,068 |
| DNA transposons (bp) | 72,935 | 199,695 | 199,936 | 226,578 | 340,490 | 255,722 | 190,480 |
| Rolling-circles (bp) | 0 | 504 | 1,414 | 704 | 0 | 1,214 | 904 |
| Unclassified (bp) | 557,170 | 1,668,439 | 1,573,557 | 1,874,045 | 1,420,719 | 1,983,403 | 2,093,113 |
| Small RNA (bp) | 5,399 | 26,790 | 30,180 | 55,375 | 428,382 | 71,525 | 103,999 |
| Satellites (bp) | 0 | 0 | 5,720 | 2,626 | 0 | 0 | 0 |
| Simple repeats (bp) | 531,479 | 551,430 | 558,219 | 555,509 | 688,325 | 599,424 | 635,054 |
| Low complexity (bp) | 130,493 | 134,560 | 132,667 | 135,193 | 153,891 | 138,519 | 145,568 |
| Total repeats (bp) | 1,711,702 | 4,470,366 | 3,972,634 | 4,906,254 | 9,754,927 | 6,886,639 | 6,200,186 |
| Non-repeat regions (bp) | 19,301,202 | 19,568,899 | 19,675,000 | 19,628,889 | 24,911,123 | 19,754,332 | 20,983,880 |

## Discussion

### Available Genome Sequencing Methods

Whole genome sequencing has revolutionized genetic studies by providing unprecedented insights into genome architecture and full genetic landscapes of an organism (Goffeau et al. 1996; Gryganskyi et al. 2023). Illumina short-read sequencing has been widely utilized and particularly for species with reference genomes, including animals (Li et al. 2009; Gerdol et al. 2020), plants (Li et al. 2020; Guo et al. 2024a), and fungi (Ahrendt et al. 2018; Lofgren et al. 2022). Its robust accuracy has been advantageous for detecting small variants and single nucleotide polymorphisms and thus been instrumental in genome-wide association studies and pangenome production across diverse organisms (Gerdol et al. 2020; Guo et al. 2024b).

However, the inherent short length of Illumina reads limits its ability to resolve repetitive elements and complex genomic regions, such as telomeric and centromeric repeats, transposable elements, and segmental duplications (Nagarajan and Pop 2013). These regions often exceed the read length, leading to ambiguous assemblies where repeats collapse into single consensus sequences.

Third-generation sequencing technologies, such as Oxford Nanopore and Pacific Biosciences (PacBio) have addressed these limitations by generating long-read solutions. Nanopore technology measures changes in electrical current as DNA molecules pass through a pore at the nanoscale; while PacBio’s SMRT sequencing relies on fluorescent signals from nucleotide incorporation (Ardui et al. 2018; Rang et al. 2018). These technologies greatly enhance genome assembly contiguity and enable the resolution of large structural variants and repetitive regions. For example, Nanopore reads have been utilized for de novo assembly of the *E. coli*, generating a complete genome and enabling an accurate reconstruction of gene order and orientation (Loman et al. 2015). However, both platforms, Oxford Nanopore and PacBio Continuous Long Reads (CLR) are prone to higher error rates due to single pass sequencing error (e.g., 5-15%), necessitating high coverage and robust bioinformatics pipelines for consensus sequence generation and error correction (Ardui et al. 2018; Rang et al. 2018). PacBio’s High-Fidelity (HiFi) sequencing combines the benefits of long-read length and high accuracy by generating consensus reads from multiple passes of circularized DNA molecules, yielding accuracies greater than 99.9% (Logsdon et al. 2020). The PacBio HiFi technology significantly improved the sequencing accuracy compared to the traditional CLR technology (Lee et al. 2023) and has been increasingly adopted in large-scale sequencing projects, such as the 1000 Fungal Genome Project, i5K Project, and Earth Biogenome Project (i5K Consortium 2013; Grigoriev et al. 2014; Lewin et al. 2018). Hybrid genome assembly approaches that integrate short-and long-read data have also gained popularity for improving genome contiguity while minimizing sequencing costs (Rech et al. 2022; Liao et al. 2023).

### De Novo Genome Sequencing of *Zancudomyces culisetae*

In this study, we compared the performance of Illumina, Oxford Nanopore, PacBio-CLR, and PacBio-HiFi sequencing technologies for de novo genome assembly of *Z. culisetae*. PacBio-HiFi emerged as the most effective platform, producing the best-quality assembly with the fewest contig numbers and the highest contiguity (Figures 2 and 3; Table 1). These findings underscore the suitability of PacBio-HiFi sequencing for generating reference-quality genomes of insect gut fungi and related organisms.

The choice of sequencing technology significantly affects genome assembly quality and computational demands. Our results show that different sequencing platforms vary in their ability to recover duplicated regions and repetitive elements. The Illumina-based assemblies exhibited smaller genome sizes (21.0-24.5 Mb), with the smallest from Illumina-only data and the largest from hybrid assembly incorporating long reads from both Nanopore and PacBio-CLR (Figures 2B and 3; Table 1). Illumina-based assemblies also contained less repetitive content (1.7-4.9 Mb), with repeat content increasing when long-read data were included (Figure 2B). Notably, the PacBio-HiFi assembly recovered significantly more retroelements (3.03 Mb) compared to the Illumina-only assembly (0.41 Mb; Table 1 and Figure 2B), consistent with the superior ability of long reads to resolve repetitive sequences, including segmental duplications and transposable elements, which tend to self-duplicate and generate large repeat-rich sections (Kwong et al. 2014; Chen et al. 2019). Interestingly, aside from the Nanopore-only assembly, all assemblies recovered similar amounts of non-repeat regions. The PacBio-HiFi assembly showed a slightly larger non-repeat portion (Figure 2B), likely due to improved recovery of duplicated genes, as indicated by a higher proportion of BUSCOs in the Complete-Duplicate category (Figure 2A).

The Nanopore assembly is the outlier with the largest genome size for both non-repeat regions (119-129% compared to other assemblies) and repeat elements (1.4-5.7x of other assemblies; Figure 2B and Table 1). It is likely that many of these repeats are false positives caused by sequencing errors. The assembler may have failed to correctly consolidate reads from the same genomic region, instead placing them in separate locations. This misassembly would artificially inflate the total genome size (Figure 3). This was also supported by the evidence that the Nanopore assembly exhibited lower contiguity and a greater number of contigs (344) than the PacBio-CLR (208) and PacBio-HiFi (26), likely reflecting sequencing errors inflated repetitive content. Although all hybrid and long-read-based assemblies recovered a higher proportion of repetitive content, the long-read only assemblies exhibit better performance in calling repetitive elements, especially the retroelements and unclassified types (Figure 2B and Table 1), which highlights the advantage of long-read sequencing in resolving complex genomic regions that are challenging to assemble with short-read data. We found that the PacBio HiFi assembly exhibits remarkable success as a unique source to generate a genome size of 28.7 Mb (Figure 4) with N50 of 2.5Mb and the largest contig of 5.2Mb, which outperformed all available genomes of insect gut fungi and provide a good reference quality genome to guide future research in this field (Wang et al. 2018; Prakash and Wang 2024).

### BUSCO Completeness and Genome Research of *Z. culisetae*

Genome completeness assessed using BUSCO revealed that the PacBio-HiFi assembly achieved the highest score (71.9%) among the assemblies, even though it still lacked approximately 20.7% of conserved genes. By comparison, the closely related soil saprobes in Kickxellomycotina (Zoopagomycota) have much higher completeness scores, such as *Coemansia reversa* (93.8%) and *Martensiomyces pterosporus* (95.1%) (Chang et al. 2015; Amses et al. 2022; Wang et al. 2023). This suggests that *Z. culisetae* has undergone significant gene loss during its evolution as an obligate insect gut symbiont when exposed to food particles, enzymes, as well as metabolites produced by the hosts and co-existing microbes (Frankel-Bricker et al. 2020; Reynolds et al. 2023). The loss is possible due to shared pathways with its hosts or co-existing microbes, a common feature of endosymbionts (Pombert et al. 2012; Sloan and Moran 2012). Over 270 million years of co-evolution with insect hosts, *Z. culisetae* and its relatives have presumably lost approximately 20% of their genes compared to their free-living relatives that share a most recent common ancestor, which has been observed in multiple studies, regardless of the sequencing methods (Wang et al. 2019; Wang et al. 2023; Prakash and Wang 2024).

As one of the first cultured microbial fungi from mosquito larvae and a globally distributed fungal symbiont, *Z. culisetae* has been instrumental in foundational discoveries on the biology of insect gut-dwelling fungi, representing an important lineage of non-Dikarya fungi.

*Zancudomyces culisetae* has pioneered multiple studies on the effects of temperature and pH on fungal development, host specificity, responses to nutritional stress, and phylogenetic relationships (Horn and Lichtwardt 1981; Lichtwardt 1986; Horn 1989; Wang et al. 2013; Van Rooy et al. 2025). However, a major obstacle to advancing our understanding of their genome biology and the molecular mechanisms underlying interactions with mosquitoes has been the lack of high-quality genome assemblies. The genomic resource of *Z. culisetae* strain HAW-14-8 presented in our study will serve as a model for advancing genomic investigations and uncovering the genome structure and hidden biology of this fungal symbiont. Notably, this strain was originally isolated from *Aedes albopictus*, an important vector of human infectious diseases, including yellow fever, dengue, and Chikungunya fever (Vega-Rúa et al. 2020; Bohers et al. 2024). The high-quality genome assembly of *Z. culisetae*, in the context of its innate association with a disease vector, provides a critical foundation for exploring host-microbe interactions and the genetic basis of fungal symbiosis in the mosquito gut environment, leading to a potential biocontrol development of insect vectors.

### Conclusion and Future Research

Our study highlights the advantages of PacBio HiFi sequencing for fungal genome assembly, offering a robust balance between sequencing accuracy, contiguity, and computational efficiency. While PacBio HiFi has higher per-base costs compared to Illumina, its ability to generate contiguous assemblies reduces the need for additional sequencing depth and complex downstream pipelines, ultimately offsetting costs. The high-quality *Z. culisetae* genome assembly provides a valuable resource for advancing research on insect gut fungi, addressing a critical gap in the field. In addition, the comprehensive comparison of sequencing technologies presented here establishes a framework for future genome projects in non-model fungal systems.

Future efforts should focus on achieving chromosome-level assemblies using ultra-long sequencing and chromatin conformation capture technologies. These advancements will further resolve complex genomic regions, including telomeres, and provide deeper insights into the evolutionary and functional adaptations of *Z. culisetae* and related fungi. By establishing standardized guidelines and high-quality reference genomes, this study lays the foundation for future genomic research on non-model organisms, particularly those with specialized ecological niches.

## Data Availability

Sequence data and the final genome assembly for *Zancudomyces culisetae* HAW-14-8 (SAMN48099576) have been deposited in the NCBI database under BioProject accession PRJNA1254182. Predicted transcripts, proteins, and all assembly files used in comparative analyses are available at https://doi.org/10.5281/zenodo.15271765.

## Acknowledgments

This research was supported by the Discovery Grants Program of the Natural Sciences and Engineering Research Council of Canada [RGPIN-2020-04293], the Discovery Launch Supplement [DGECR-2020-00154], and Connaught New Researcher Award [NR-2021-22-514711]. Computations were performed on the Niagara supercomputer at the SciNet HPC Consortium. SciNet is funded by Innovation, Science and Economic Development Canada; the Digital Research Alliance of Canada; the Ontario Research Fund: Research Excellence; and the University of Toronto. The authors thank Tara Paton for valuable discussions and sequencing support.

